# Engineering of the Endogenous *HBD* promoter increases HbA2

**DOI:** 10.1101/2022.12.19.521003

**Authors:** Mandy Y. Boontanrart, Elia Mächler, Simone Ponta, Jan C. Nelis, Viviana G. Preiano, Jacob E. Corn

## Abstract

The β-hemoglobinopathies, such as sickle cell disease and β-thalassemia, are one of the most common genetic diseases worldwide and are caused by mutations affecting the structure or production of β-globin subunits in adult hemoglobin. Many gene editing efforts to treat the β-hemoglobinopathies attempt to correct β-globin mutations or increase γ-globin for fetal hemoglobin production. δ-globin, the subunit of adult hemoglobin A2, has high homology to β-globin and is already pan-cellularly expressed at low levels in adult red blood cells. However, upregulation of δ-globin is a relatively unexplored avenue to increase the amount of functional hemoglobin. Here, we use CRISPR-Cas9 to repair non-functional transcriptional elements in the endogenous promoter region of δ-globin to increase overall expression of adult hemoglobin 2 (HbA2). We find that insertion of a KLF1 site alone is insufficient to upregulate δ-globin. Instead, multiple transcription factor elements are necessary for robust upregulation of δ-globin from the endogenous locus. Promoter edited HUDEP-2 immortalized erythroid progenitor cells exhibit striking increases of *HBD* transcript, from less than 5% to over 20% of total β-like globins. Edited CD34+ hematopoietic stem and progenitors (HSPCs) differentiated to primary human erythroblasts express up to 35% *HBD*. These findings add mechanistic insight to globin gene regulation and offer a new therapeutic avenue to treat β-hemoglobinopathies.

## Introduction

Red blood cells, also known as erythrocytes, are packed with hemoglobin and circulate the body to supply all organs with oxygen. Hemoglobin is a hetero-tetrameric protein made up of two α-like and two β-like subunits. Hemoglobin A1 (HbA1) accounts for approximately 97% of the hemoglobin expressed in adults and is composed of two α-globin subunits (*HBA*) and two β-globin (*HBB*) subunits. Hemoglobin A2 (HbA2) accounts for the remaining 2-3% of hemoglobin expressed in adults and is composed of two α-globin subunits and two δ-globin (*HBD*) subunits^1^.

The β-hemoglobinopathies, such as sickle cell disease (SCD) and β-thalassemia, are caused by mutations in *HBB* that effect the structure or expression of β-globin. The clinical hallmarks include hemolytic anemia and vaso-occlusion, which can lead to acute and chronic pain and organ damage. Clinical management is limited to frequent blood transfusions and life-long treatment of anemia and pain crises. The only curative approach is allogeneic stem cell transplantation, which is dependent upon HLA-identical donor availability^2^. Fetal hemoglobin (HbF), which is the predominant hemoglobin expressed before birth, has anti-sickling properties and its re-expression is frequently pursued as a treatment for β-hemoglobinopathies^3^. While increasing HbF has shown to be clinically effective to combat SCD, studies have also validated the *in vitro* and *in vivo* anti-sickling abilities of δ-globin using a humanized mouse model of SCD^4–7^.

Increased HbA2 expression has some potential advantages over HbF that suggests it could provide an alternative avenue for to ameliorate β-hemoglobinopathies. While HbF expression is completely silenced after birth, HbA2 is weakly transcriptionally active and expressed pancellularly in all adult red blood cells ^8,9^ Additionally, δ-globin shares 93% homology to β-globin, suggesting that HbA2 may be a better replacement relative to HbF, whose γ-globin subunit shares 73% homology to β-globin. Finally, HbF is known to bind oxygen more tightly than HbA1, an evolutionary advantage for the fetus to draw oxygen from the maternal blood source, while HbA2 has a similar oxygen-binding capacity as HbA1^10,11^. On-going clinical trials re-express HbF to extremely high levels not commonly seen even in individuals with Hereditary Persistence of Fetal Hemoglobin^12^. The effect of extreme maternal HbF re-expression during pregnancy is currently unknown.

The genes for β-globin (*HBB*) and δ-globin (*HBD*) are located in the β-like globin cluster and regulated by the same control region. The β-like globin cluster is located on chromosome 11, and harbors the five β-like genes: *HBB* (β-globin gene), *HBD* (δ-globin gene), *HBG1* and *HBG2* (γ-globin genes), and HBE (ε-globin gene). β-globin and δ-globin both comprise 147 amino acids and differ at only 10 positions^13^. The extreme difference in expression levels between these two globins is not due to protein instability or differences in translation, but instead results from a lower transcription rate^1^. The globin genes are arranged in order of their expression during development and regulated by contact to a distal enhancer called the Locus Control Region that contains five active DNase Hypersensitivity Sites^14^. A comparative genomics study has shown that, compared to the *HBB* promoter, the *HBD* promoter has mutations in multiple transcriptional elements including a KLF1, CP1, β-DRF and TFIIB binding site (**Fig. 1A**)^15^.

**Figure 1.**
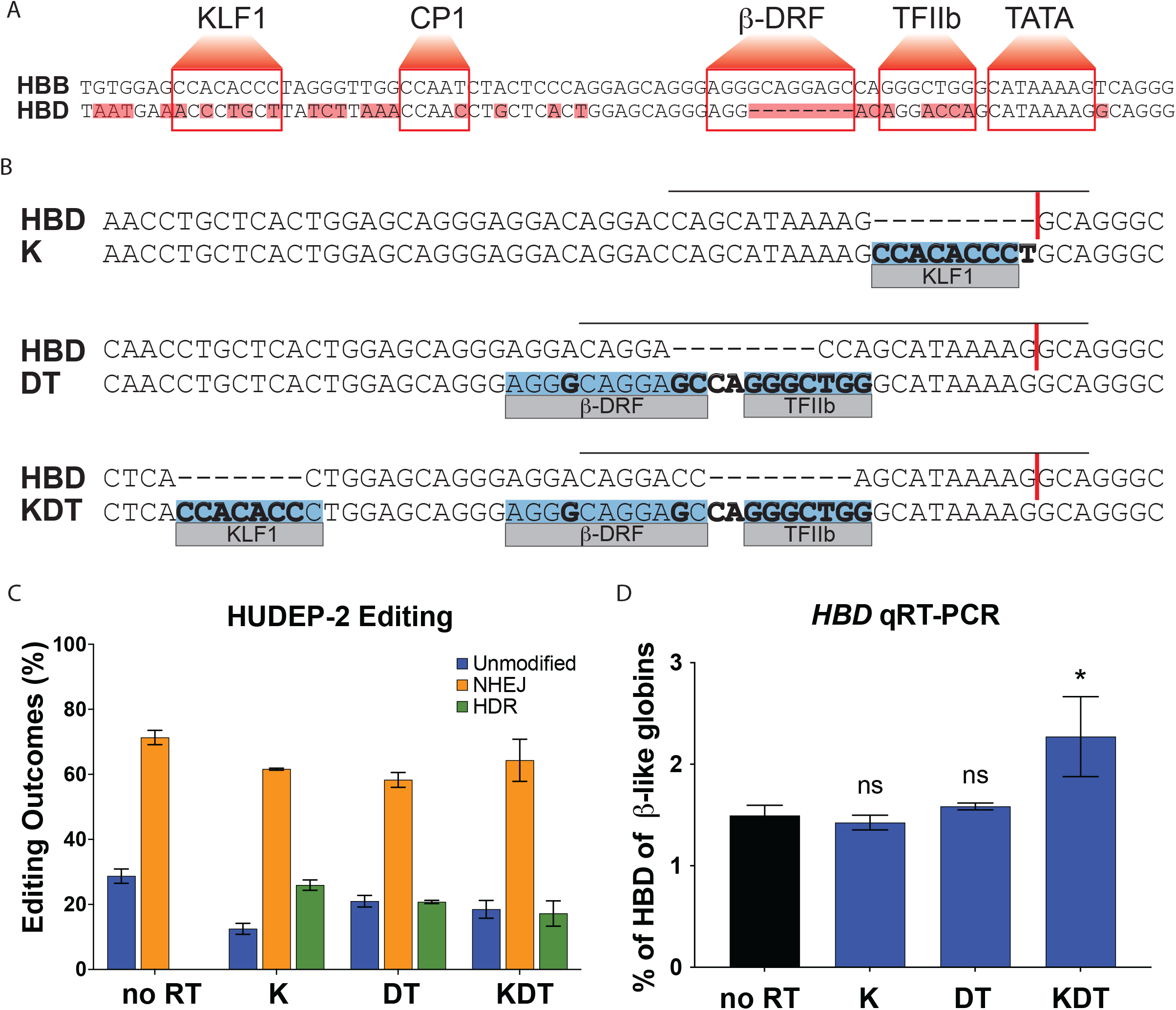
Targeting and design of the endogenous *HBD* promoter. A) Alignment of the *HBB* and *HBD* promoter sequences. Transcription factor binding sequences for KLF1, CP1, β-DRF, TFIIb, and TATA are shown in boxes and base pair mismatches between *HBB* and *HBD* are highlighted in red. B) The repair template designs for insertions of KLF1 (K), β-DRF and TFIIb (DT), and KLF1, β-DRF, and TFIIb (KDT) directly compared to the *HBD* promoter. The inserted transcription factor binding sequences are highlighted in blue. Any base pair changes are in bold. The gRNA is indicated by a black horizontal line and the cut site is indicated by a red vertical line. C) HUDEP-2 editing efficiencies showing percentages of unmodified, NHEJ, or HDR alleles. Conditions tested were Cas9 and sgRNA RNP with no repair template (no RT), K, DT, and KDT repair templates. This experiment was performed three times and the data is presented as mean ± SD of three biological replicates. D) qRT-PCR of *HBD* after pooled editing of HUDEP-2 cells with no RT, K, DT, and KDT and 5 days of differentiation. Data is plotted as % of all β-like globins (*HBB, HBG1/2, HBD*). The three biological replicates from the editing experiment in C) were each differentiated and the data is presented as mean ± SD of three biological replicates. P value indicates paired, two-tailed student *t* test (ns, non-significant; *, P ≤ 0.05; **, P ≤ 0.01).

Previous studies using transgenic approaches have shown that inclusion of a KLF1 motif in the *HBD* promoter can drive exogenous expression of δ-globin ^6,16,17^. However, these studies do not reflect the complex chromosomal context and extensive epigenetic regulation at the β-like globin cluster. Due to the large size of the β-like globin locus, transgenic studies have mostly included only a subset of the genes of the β-like globin locus and a minimal region of the LCR. They therefore do not necessarily predict the biological outcomes of perturbations at the native β-globin locus^18^.

The expression of globin genes is a tightly regulated development process. There are currently no drugs or therapeutic approaches to increase HbA2 for the β-hemoglobinopathies. Using CRISPR-Cas9 genome editing, we used homology directed repair at the endogenous *HBD* promoter to engineer the transcriptional elements present in *HBB*. We find that insertion of single transcriptional elements to the endogenous promoter is insufficient for δ-globin upregulation. However, insertion of KLF1, β-DRF, and TFIIB motifs drive high expression of δ-globin from the endogenous locus in HUDEP-2 cells and primary erythroblasts. This leads to reconstitution of high levels of HbA2, over 10-fold increase compared to WT unedited cells. Our work adds mechanistic insight to the globin gene regulation at the β-like globin cluster and suggests a potential therapeutic avenue to upregulate HbA2 for the β-hemoglobinopathies.

## Results

### Targeting the endogenous *HBD* promoter to introduce functional *HBB* promoter elements

We aligned the promoter sequences of *HBB* and *HBD* to identify transcriptional elements missing in the *HBD* promoter (**Fig 1a**). This highlighted multiple mutations and deletions in the KLF1, CP1, β-DRF, and TFIIB binding motifs. To re-engineer the endogenous *HBD* promoter, we employed CRISPR-Cas9 induced homology directed repair (HDR) gene editing ^19,20^.

We designed and tested three sgRNAs targeting the *HBD* promoter **(Fig S1A)** and three HDR templates that would incorporate base pairs needed to complete the transcriptional element motifs (**Fig 1b**). The HDR templates were designed as single stranded oligo donor nucleotides (ssODN) to either insert a KLF1 (K) sequence, a β-DRF and TFIIB (DT) sequence, or all three elements (KDT). The β-DRF and TFIIB motifs were designed into a single template because they are separated by only 2 base pairs. A previous transgenic study showed that the CP-1 site plays a minor role in *HBB* transcription, and was therefore omitted from our designs^21^. We maintained the ordering between motifs to mimic the *HBB* promoter and paid particular attention to limiting the number of mutated base pairs to increase the likelihood of successful HDR editing.

We performed editing in HUDEP-2 cells, an immortalized cell line capable of differentiating into hemoglobin-producing erythroid cells ^22^, using Cas9 ribonucleoprotein (RNP) and an ssODN. After optimizing conditions and testing multiple sgRNAs (**Table S1**), we found NHEJ rates ranging from 56 – 74% and HDR efficiencies from 13 – 27% as measured by next generation sequencing (NGS) (**Fig 1c**). The edited pools were differentiated to erythroblasts and qRT-PCR measurements were taken to assess the effects on *HBD* expression (**Fig 1d**). Despite roughly equal HDR rates between all promoter designs, we observed a statistically significant increase in *HBD* only for the KDT design with three elements.

### Homozygous knock-in of KLF1, TFIIB, and β-DRF leads to robust increase of *HBD* in HUDEP-2 cells

Heterogeneously edited pools of cells, containing a mixture of alleles, can mask large effects at a clonal level. We isolated heterozygous and homozygous HDR clones to more accurately assess the effect of each motif edit on *HBD* expression. We obtained at least one heterozygote and homozygote for each edit, as verified by amplicon NGS sequencing of the *HBD* promoter (**Fig S1**). We used ChIP-qPCR to test whether the inserted motifs were functional in recruiting KLF1 and RNA Pol II to the *HBD* promoter, comparing to the *HBB* promoter and *VEGFA* as a positive and negative control, respectively. Due to the unavailability of a TFIIB antibody suitable for ChIP, we performed ChIP for RNA Pol II, which is a binding partner of TFIIB in the pre-initiation complex (**Fig 2a**). The β-DRF motif has been shown to be important for high transcription of *HBB*, but its *bona fide* binding factor has not yet been identified^23^. In the homozygous clones K and KDT, which harbor the edited KLF1 sequence, we observed increased binding of KLF1 to the *HBD* promoter relative to unedited WT HUDEP-2 cells. However, we found RNA Pol II binding at the *HBD* promoter binding only in the homozygous KDT clone. The homozygous DT clone, which harbors a TFIIB site, did not show significant binding of RNA Pol II. Taken together, these data indicate that an intact KLF1 site is sufficient to recruit KLF1, but all three transcription factor binding sites are necessary to recruit the transcriptional machinery to the *HBD* promoter ^24^.

**Figure 2.**
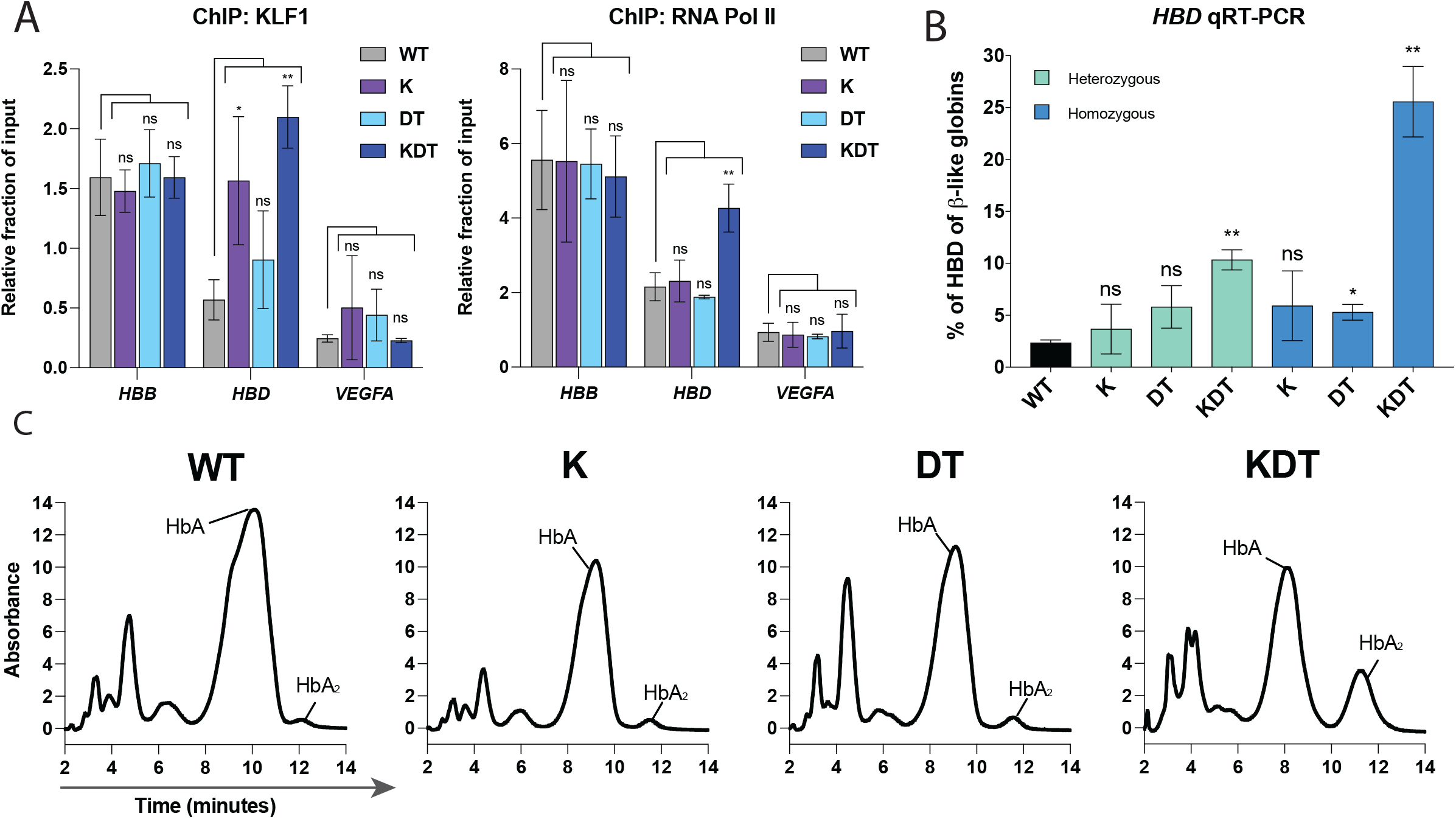
Characterization of HUDEP-2 clones with heterozygous or homozygous knock-ins of KLF1, β-DRF, and TFIIb sequences. A) ChIP-qPCR of KLF1 and RNA Pol II performed on WT HUDEP-2 cells and homozygous clones of K, DT, and KDT. Data is shown as relative fraction of input and normalized to *SP1*. The genes targeted are *HBD, HBB*, and *VEGFA* as a negative control. Cells were grown and harvested separately and on different days for each biological replicate. The data is presented as mean ± SD of three biological replicates. P value indicates paired, two-tailed student *t* test (ns, non-significant; *, P ≤ 0.05; **, P ≤ 0.01). B) qRT-PCR of *HBD* of HUDEP-2 heterozygous and homozygous clones with K, DT, and KDT knock-in and 5 days of differentiation. Data is plotted as % of β-like globins (*HBB, HBG1/2, HBD*). Each clone was differentiated on three separate days and the data is presented as mean ± SD of three biological replicates. P value indicates paired, two-tailed student *t* test (ns, non-significant; *, P ≤ 0.05; **, P ≤ 0.01). C) HPLC of of HUDEP-2 homozygous clones with K, DT, and KDT knock-in and 5 days of differentiation. Hemoglobin A (HbA) and Hemoglobin A2 (HbA2) peaks are annotated.

We next performed qRT-PCR on all differentiated heterozygous and homozygous clones to look at percentage of *HBD* produced compared to total β-like globins (**Fig 2b**). *HBB* expression was unaffected by the K, DT, and KDT knock-ins **(Fig S2).** Like the pooled editing results, we found that clones with single knock-ins of KLF1 or β-DRF and TFIIB motifs alone did not show increased *HBD*. However, we observed a striking increase in *HBD* for KDT clones harboring all three motifs. The heterozygous clone showed *HBD* increases to 10% of total β-like globins. The homozygous clone showed increases in *HBD* from less than 5% to over 20% of β-like globins.

Next, hemoglobin protein levels were measured using high pressure liquid chromatography (HPLC) for the differentiated homozygous clones (**Fig 2c**). Peaks were assigned to hemoglobin complexes based on previous work that performed mass-spectrometry on each globin peak fraction ^25^. We found that the transcript-level results are further supported at the protein level. The KLF1 and the β-DRF and TFIIB clones did not show any measurable increase in HbA2, while the KDT homozygous showed a large increase in HbA2.

### Endogenous editing of the *HBD* promoter increases *HBD* in CD34+ derived erythroblasts

To test the effects of *HBD* promoter engineering in a more clinically relevant cell type for a potential *ex vivo* therapy to treat β-hemoglobinopathies, we edited human CD34+ derived erythroblasts. We used CRISPR-Cas9 to perform pooled knock-in at the *HBD* promoter in mobilized peripheral blood CD34+ human hematopoietic stem and progenitor cells (HSPCs) with the K, DT, and KDT RNP and ssODNs. We found HDR rates ranging from 18 – 25% for all promoter knock-ins (**Fig 3a**). We differentiated the edited HSPC pools and performed qRT-PCR to measure *HBD* expression levels. As in HUDEP-2 cells, there was no statistically significant increase in *HBD* levels for K and DT, and a slight increase in two different donors in the KDT knock-in condition (**Fig 3b**).

**Figure 3.**
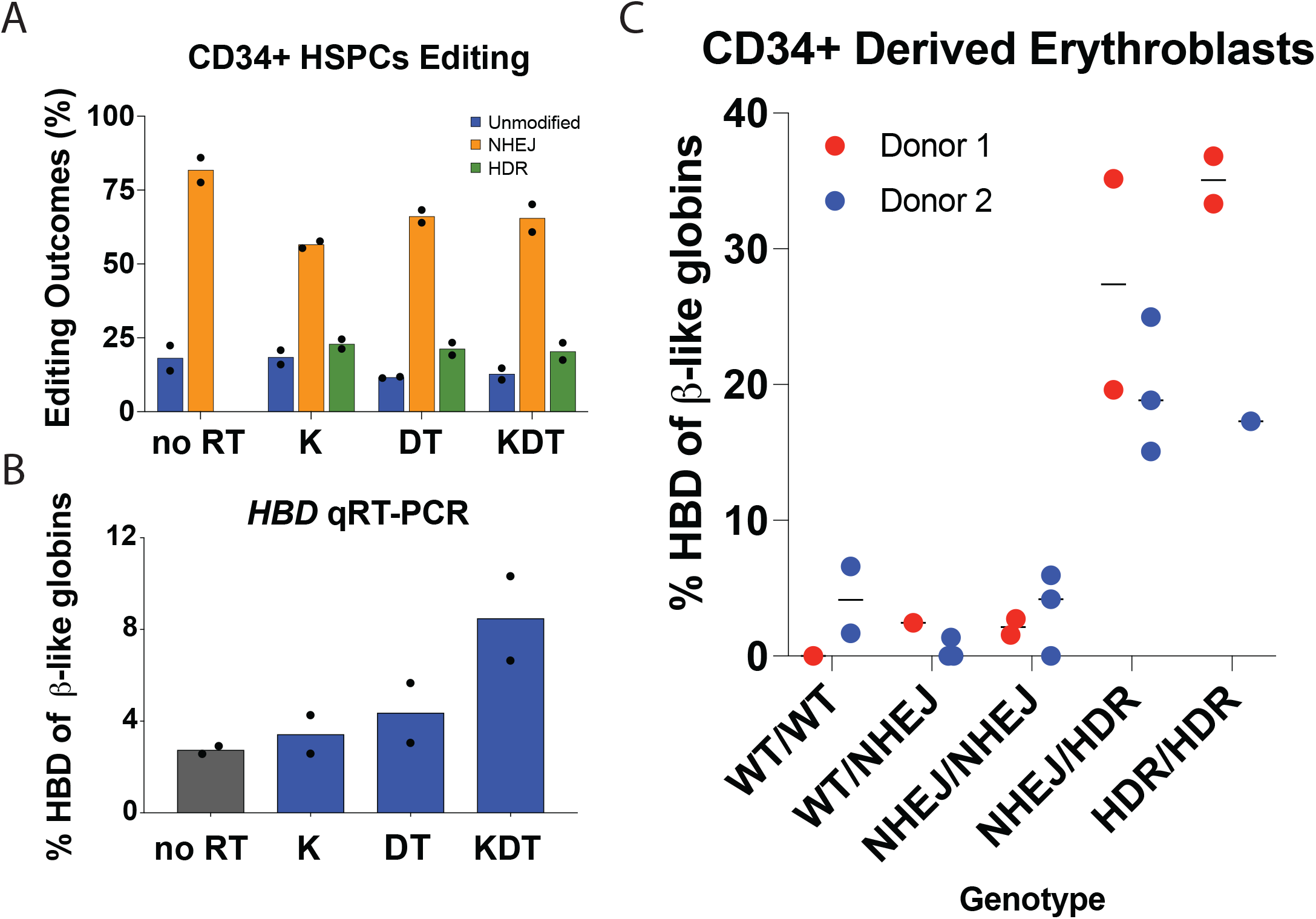
Endogenous editing of the *HBD* promoter in HSPCs. A) Editing efficiencies showing percentages of unmodified, NHEJ, or HDR alleles. Conditions tested were Cas9 and sgRNA RNP with no repair template (no RT), K, DT, and KDT repair templates. The data is presented as two independent editing experiments with two different donor samples. B) qRT-PCR of *HBD* after pooled editing of HSPCs with no RT, K, DT, and KDT. Cells were expanded in erythroid expansion conditions and differentiated for 5 days. Data is plotted as % of all β-like globins (*HBB, HBG1/2, HBD*). The data is presented as two independent editing experiments with two different donor samples. C) qRT-PCR of *HBD* of clonal erythroblast populations after 5 days of differentiation. Genotypes were determined by NGS. Data is plotted as % of β-like globins (*HBB, HBG1/2, HBD*). The data is presented as two independent editing experiments with two different donor samples and a dot denotes individual clonal populations.

To test the effect of KDT editing on a single-cell level, we isolated and grew colonies from HSPCs under erythroid expansion conditions. Each colony was genotyped and their alleles classified as either unmodified, NHEJ, or HDR (**Fig S3A**). These CD34+ derived clonal erythroblasts were differentiated and qRT-PCR was performed to determine *HBD* expression relative to total β-like globins **(Fig 3c**). *HBB* transcript levels remain similar between unedited or NHEJ clones and KDT knock-in clones (**Fig S3B**). We observed that colonies with unedited or all NHEJ alleles had an average expression of lower than 3% *HBD* of total β-like globins. Heterozygous KDT knock-ins from multiple donors increased *HBD* expression dramatically, between 15 – 35% of total β-like globins in heterozygous knock-ins and between 17 – 37% of total β-like globins in homozygous knock-ins.

## Discussion

Current gene editing efforts to treat the β-hemoglobinopathies include correcting individual *HBB* mutations^26^, a method which would be limited to specific types of patient mutations, or increasing fetal hemoglobin^3^, a gene normally silenced after birth and with different oxygen-binding capacities than HbA1. In this study, we describe a path to upregulating HbA2, which shares high similarity to HbA1 and is applicable to all β-hemoglobinopathy disease mutations.

Previous transgenic studies have shown that insertion of KLF1 alone to the *HBD* promoter sequence is sufficient to drive expression of *HBD* ^6,16^. However, these studies do not reflect the complex chromatin context and regulation at the β-like globin locus. We find that endogenous insertion of a KLF1 motif alone is insufficient to drive *HBD* expression. Instead, insertion of three motifs, KLF1, β-DRF, and TFIIB, is necessary to recruit RNA Pol II and induce high expression of *HBD*. In CD34+ derived erythroblasts, homozygous knock-in of KLF1, β-DRF, and TFIIB leads to increases in *HBD* up to 35% of total β-like globins. To our knowledge, this is the first genomic editing of the *HBD* promoter that results in increased HbA2. Bulk editing of *HBD* through insertion of multiple transcriptional elements will rely on high levels of HDR. Increasing HDR outcomes might be achieved in CD34+ cells by a variety of methods such as controlled cell-cycling or modulation of DNA repair factors ^27,28,29,30^.

Our promoter engineering approach dramatically increases *HBD* levels, but it is not yet clear whether *HBD* can be increased to the same levels as *HBB*. Using ChIP, we observed somewhat lower RNA Pol II binding at the *HBD* promoter than at the *HBB* promoter in the homozygous KDT clone. This suggests that further engineering of the *HBD* promoter could possibly increase *HBD* transcription. The β-globin locus is tightly regulated by developmental epigenetics and erythroid-specific transcription factors. Further work might explore the insertion of more erythroid-specific or synthetic transcription factor binding sites in attempts to further increase in *HBD*.

It is estimated that roughly 30% of beneficial hemoglobin (10 pg per cell) is sufficient to ameliorate β-hemoglobinopathy symptoms^9^. In our experiments, heterozygous and homozygous knock-in of KDT in CD34+ erythroblasts led to 15 – 37% *HBD* relative to total β-like globins. Interestingly, in edited CD34+ erythroblasts, we observed that heterozygous and homozygous KDT populations expressed similar increases in *HBD*. Further studies should investigate whether heterozygous knock-in of KDT in β-hemoglobinopathy cells is sufficient to ameliorate disease phenotypes. For example, one could knock-in KDT to the *HBD* promoter of SCD patient HSPCs and perform HPLC or microscopy assays to determine the anti-sickling effects of a heterozygous or homozygous knock-in. If heterozygous knock-in of KDT is sufficient, one would see a decrease in sickle hemoglobin produced, or a decreased percentage of sickled red blood cells.

In summary, our presented study provides a novel strategy to increase levels of HbA2 from the endogenous *HBD* locus that could potentially be applicable as an *ex-vivo* gene therapy. In pre-clinical studies, it will be important to quantitatively investigate the amount of *HBD* that is optimal for improving the function and health of patient red blood cells and explore the overall potential and safety of increasing HbA2 as a therapeutic option for the β-hemoglobinopathies.

## Acknowledgements

We thank laboratory members for helpful discussions and support; D. Fercher and M. Zenobi-Wong for help with HPLC; R. Kurita and Y. Nakamura for their contribution of HUDEP-2 cells; Genome Engineering Measurement Lab and Functional Genomics Center Zurich (ETH Zurich/University of Zurich) for help in running NGS samples. J.E.C. is supported by the NOMIS Foundation and the Lotte and Adolf Hotz-Sprenger Stiftung. M.Y.B is supported by ETH Foundation’s ETH Pioneer Fellowship and the SNSF BRIDGE Foundation.

## Author Contributions

M.Y.B. designed the experiments.

M.Y.B., E.M., S.P., J.C.N., V.G.P. performed experiments.

M.Y.B., E.M., S.P., J.C.N. analyzed data.

M.Y.B. and J.E.C. wrote the manuscript.

All authors contributed feedback for the manuscript.

## Figure Legends

The data is presented as mean ± SD of three biological replicates. P value indicates paired, twotailed student *t* test (ns, non-significant; *, P ≤ 0.05; **, P ≤ 0.01).

## Supplementary Figure Legend

**Figure S1.**
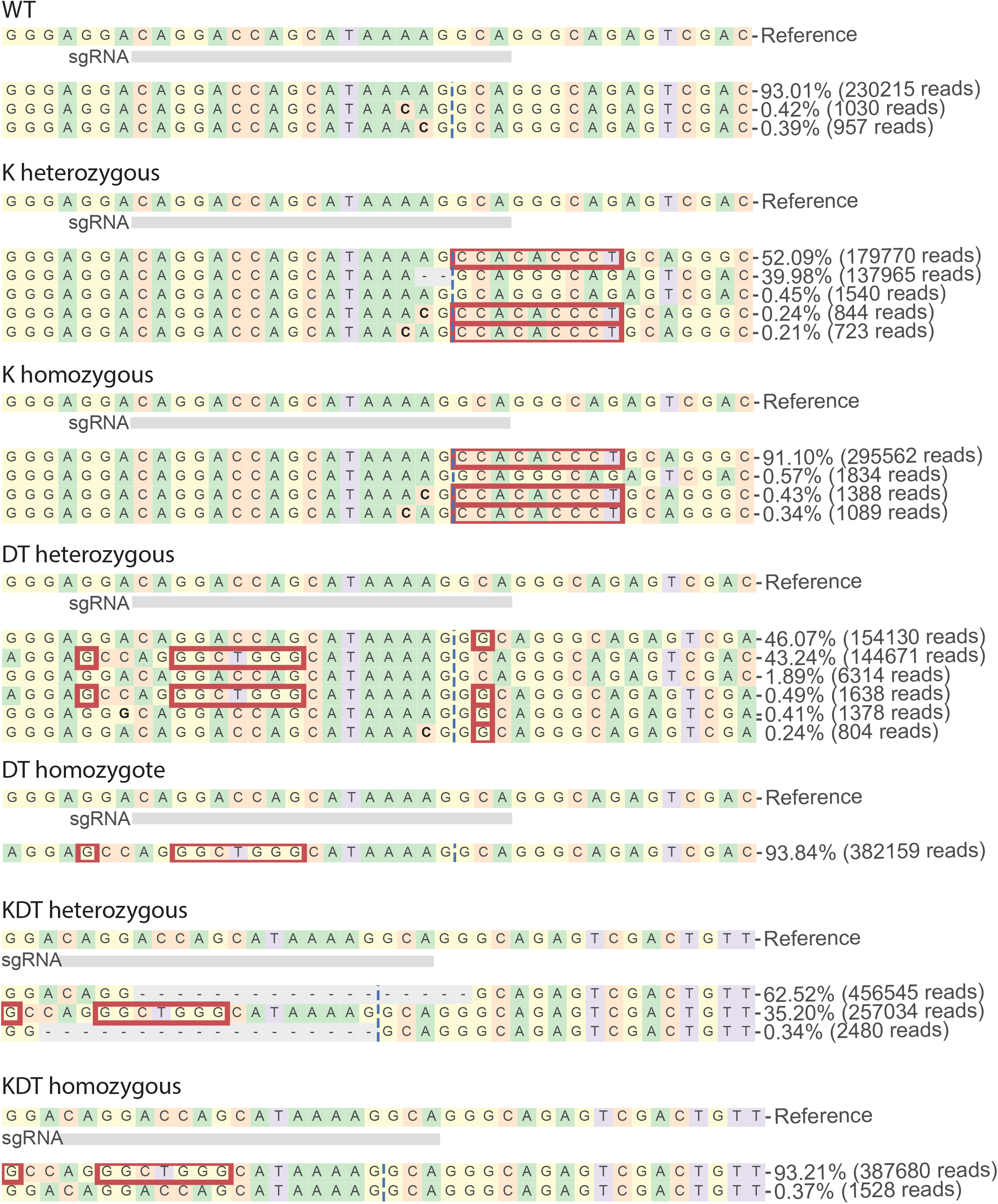
Genotypes of HUDEP-2 clones with heterozygous or homozygous knock-ins of KLF1, β-DRF, and TFIIb sequences Clonal HUDEP-2 cell lines of K, DT, and KDT promoter knock-ins were generated and genotypes using NGS and CRISPResso analysis. Red boxes denote insertions.

**Figure S2.**
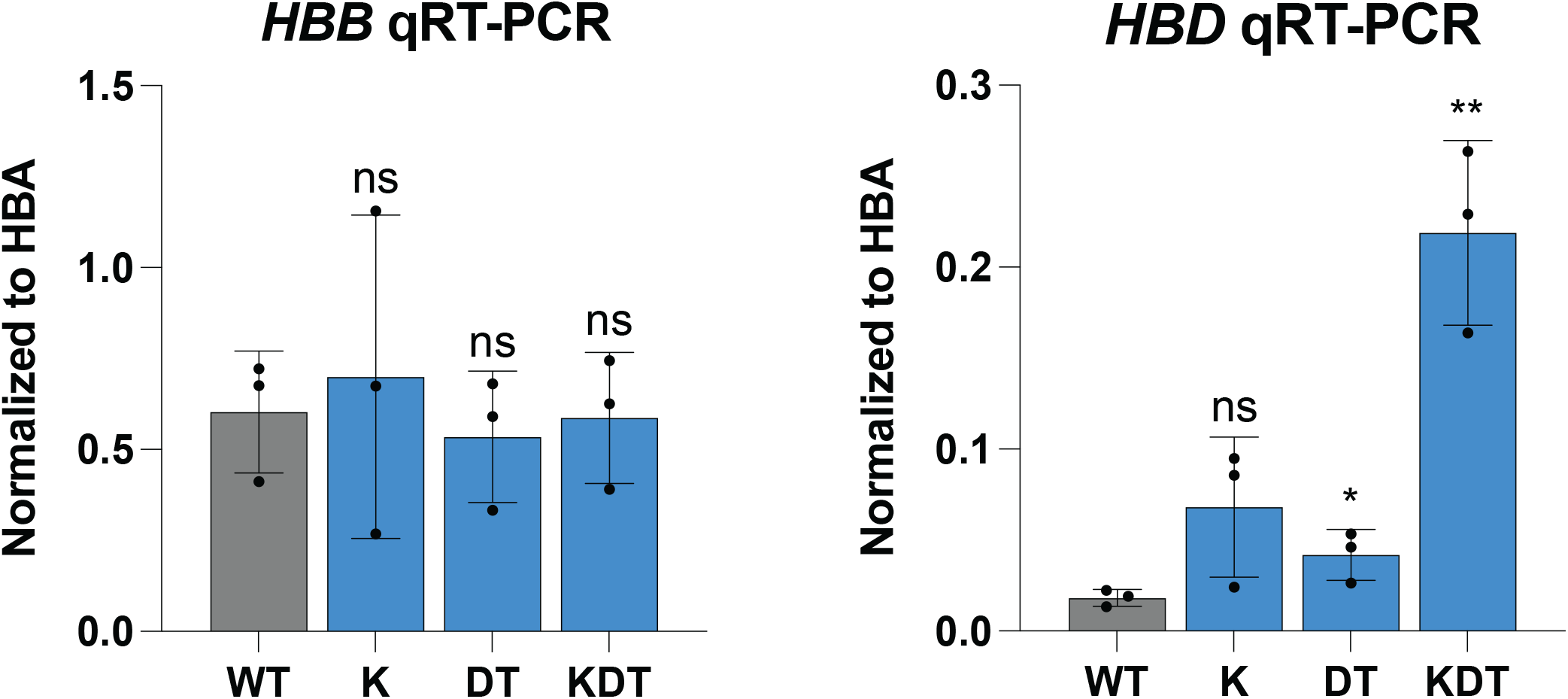
qRT-PCR of HUDEP-2 clones with heterozygous or homozygous knock-ins of KLF1, β-DRF, and TFIIb sequences qRT-PCR of *HBD* of HUDEP-2 heterozygous and homozygous clones with K, DT, and KDT knock-in and 5 days of differentiation. Data is plotted as normalized to *HBA*. Each clone was differentiated on three separate days and the data is presented as mean ± SD of three biological replicates. P value indicates paired, two-tailed student *t* test (ns, non-significant; *, P ≤ 0.05; **, P ≤ 0.01).

**Figure S3.**
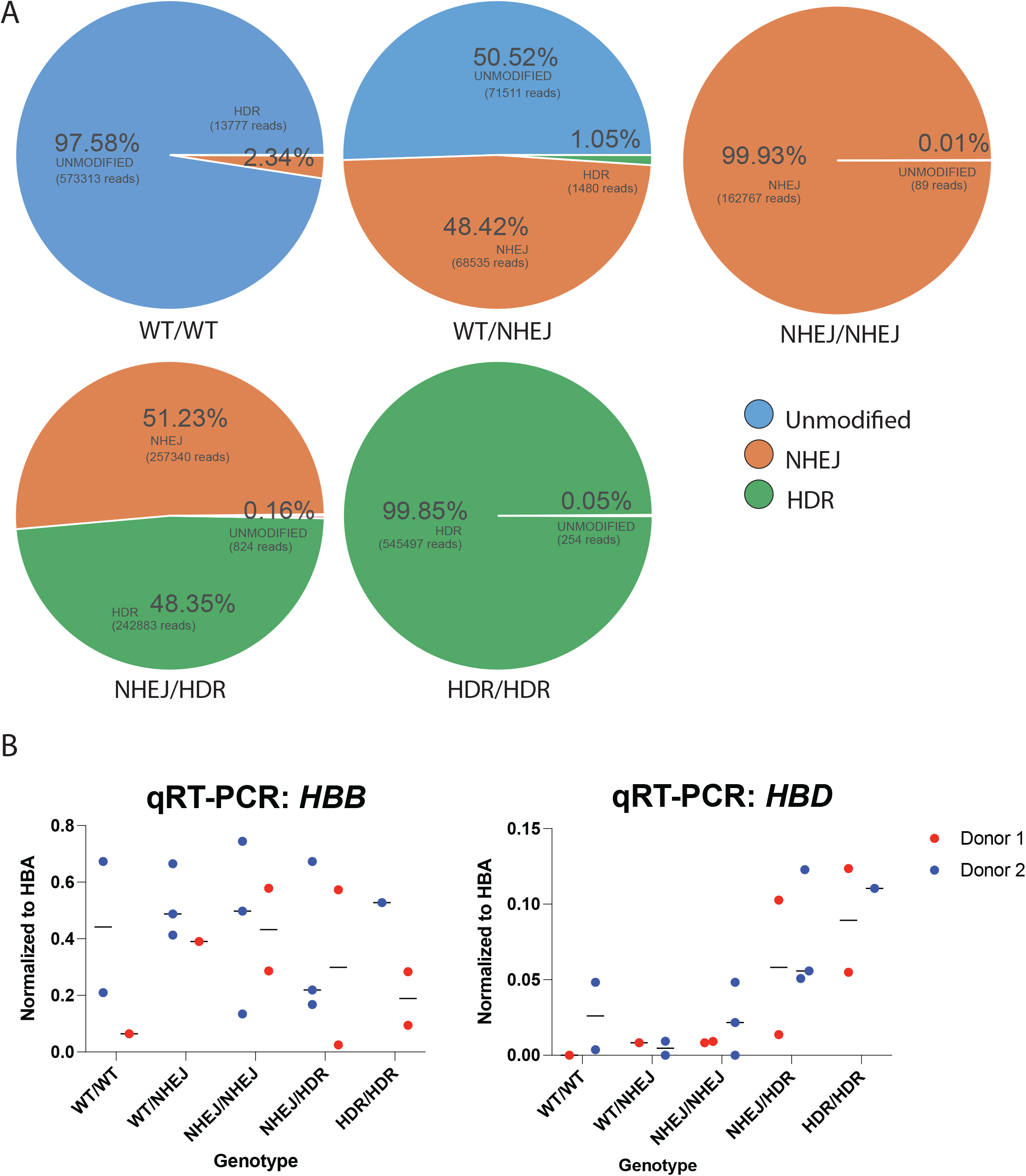
Characterization of *HBD* promoter edited CD34+ HSPCs. A) An example subset of genotypes of WT/WT, WT/NHEJ, NHEJ/NHEJ, NHEJ/HDR, and HDR/HDR clonal erythroblasts. B) qRT-PCR of *HBD* of clonal erythroblast populations after 5 days of differentiation. Genotypes were determined by NGS. Data is plotted as normalized to *HBA*. The data is presented as two independent editing experiments with two different donor samples and a dot denotes individual clonal populations.

## Methods and Materials

### Cas9 RNP Nucleofection

Cas9 RNP was performed as described previously (Lingeman et al., 2017). Briefly, either IVT guides are purified or chemically protected guides were ordered from Synthego and complexed with purified Cas9-NLS protein. The nucleofection was performed using Lonza 4D-Nucleofector and using the P3 Primary Cell 96-well NucleofectorTM Kit (V4SP-3096) following manufacturer’s instructions. The HUDEP-2 nucleofector code used was DD-100 and for primary HSPCs ER-100.

### IVT sgRNA

Guide RNAs (Table S1) were in vitro transcribed as described previously (Lingeman et al., 2017). Briefly, guide sequences were ordered as oligonucleotides and formed into duplexes using a PCR thermocycler. The DNA template was transcribed to RNA using HiScribe ™ T7 High Yield RNA Synthesis Kit (E2040S) following manufacturer protocol. The resulting RNA was purified using RNeasy Mini kit (74104) and Rnase-Free DnaseI Kit (79254).

### High Pressure Liquid Chromatography and Mass Spectrometry

HUDEP-2 cells or HSPCs were differentiated and harvested for lysis in hemolysate reagent containing 0.005M EDTA and 0.07% KCN at 10,000 cells per microliter. The lysis was incubated at room temperature for ten minutes and then centrifuged at max speed for 5 minutes. The supernatant was collected and run on Agilent 1260 Infinity II using a PolyCAT A column from PolyLC, 35×4.6mm (3μm;1500Å) Serial# B19916E. The following Buffer compositions were used: Mobile Phase A: 20mM Bis-tris, 2mM NaCN pH 6.8 and Mobile Phase B: 20mM Bis-tris, 2mM NaCN, 200mM NaCl, pH 6.9. The following flow settings were used: Gradient: 0-8’ 2-25% Phase B, 8-18’ 25-100% Phase B, 18-23’ 100-2% Mobile Phase B using a Flow Rate: 1.5mL/min and measuring detection of 415nm Diode Array.

### HUDEP-2 cell culture and differentiation

All cell culture was performed at 37°C in a humidified atmosphere containing 5% CO2. HUDEP-2 cells (RRID: CVCL_VI06), obtained from the Riken Institute^22^, were cultured in a base medium of SFEM (Stemcell Technologies 9650) containing to a final concentration of dexamethasone 1uM (Sigma D4902-100MG), doxycycline 1ug/ml (Sigma D9891-1G), human stem cell factor 50ng/ml (PeproTech 300-07), erythropoietin 50ng/ml (Peprotech 100-64), and penstrept 1%. Cells were cultured at a density of 2e5 – 1e6 cells/ml. For differentiation, HUDEP-2 cells are centrifuged at 500g for 5 minutes, media is removed and replaced with differentiation media. Differentiation media consists of a base media of IMDM+Glutamax (ThermoFisher 31980030) containing to a final concentration human serum 5% (Sigma H4522-100mL), heparin 2IU/ml (Sigma H3149-25KU), insulin 10ug/ml (Sigma I2643-25mg), erythropoietin 50ng/ml (Peprotech 100-64), holo-transferrin 500ug/ml (Sigma T0665-100mg), mifepristone 1uM (Sigma M8046-100MG), and doxycyline 1ug/ml (Sigma D9891-1G). Cells are differentiated for 5 days and then harvested for analysis.

### mPB-HSPCs cell culture and differentiation

For editing for human CD34+ cells, CD34+ mobilized peripheral blood HSPCs were thawed and cultured in SFEM containing CC110 supplement (Stemcell Technologies 02697) for 2 days. CD34+ cells were then electroporated and transferred into erythroid expansion media containing SFEM and erythroid expansion supplement (Stemcell Technologies 02692) for 7 days and cultured at a density of 2e5-1e6 cells/ml. The resulting early erythroblasts were transferred to differentiation media containing SFEM with 50ng/ml erythropoietin, 3% normal human serum, and 1 μM mifepristone. The resulting late erythroblasts were harvested for analysis after 5 days of differentiation. For generation of clonal erythroblasts, CD34+ cells were then electroporated and transferred into erythroid expansion media containing SFEM and erythroid expansion supplement (Stemcell Technologies 02692) for 4 days. The early erythroblasts were then single cell seeded and cultured for 7 days. After 7 days, the cells are transferred into differentiation media containing SFEM with 50ng/ml erythropoietin, 3% normal human serum, and 1 μM mifepristone. The resulting late erythroblasts were harvested for analysis after 5 days of differentiation.

### qRT-PCR

RNA was harvested from cells using Qiagen RNeasy Mini Kit and Rnase-Free DnaseI Kit following manufacturer’s instructions. RNA was reverse transcribed to cDNA using Iscript™ Reverse Transcription Supermix (BioRad) and qRT-PCR reactions were set up using SsoAdvanced Universal SYBR Green or SsoFast™ EvaGreen Supermix (BioRad). Reactions were run on the StepOne Plus Real-Time PCR System (Applied Biosystems) or the QuantStudio 6 Flex (Thermo Fisher). Samples were analyzed using a two-step amplification and melt curves were obtained after 40 cycles. The Ct values for genes of interest were normalized to GAPDH, and expressions of genes are represented as 2-[ΔCt] or 2-[ΔΔCt] for fold change over control condition. All primers used for qRT-PCR are listed in Table S1.

### ChIP-qPCR

ChIP was performed as done previously^25^. Briefly, 10 million cells per sample were harvested and cross-linked in 1% Formaldehyde. Cross-linking was quenched with the addition of 1.5M glycine. Samples were then lysed for 10 minutes at 4°C in 50 mM Hepes-KOH, pH 7.5; 140 mM NaCl; 1 mM EDTA; 10% glycerol; 0.5% NP-40 or Igepal CA-630; 0.25% Triton X-100. Cells were then centrifuged at 1500g for 3 minutes and the supernatant was discarded. The pellet was resuspended in 10 mM Tris–HCl, pH8.0; 200 mM NaCl; 1 mM EDTA; 0.5 mM EGTA and incubated for 5 minutes at 4°C. The cells were then centrifuged at 1500g for 3 minutes and the supernatant was discarded. The pellet was resuspended in 10 mM Tris–HCl, pH 8; 100 mM NaCl; 1 mM EDTA; 0.5 mM EGTA; 0.1% Na–Deoxycholate; 0.5% N-lauroylsarcosine and sonicated using the Covaris S220 following manufacturer’s instructions. Protein G beads (ThermoFisher) were complexed with antibody and the antibody-bead complexes were incubated with cell lysates at 4C overnight with rotation. The antibodies used were mouse anti-RNA Pol II (Diagenode C15100055-100 RRID: AB_2750842) and goat anti-KLF1 (Origene TA305808). The beads were retrieved using a magnetic stand and rinsed with RIPA buffer. Elution buffer containing 50 mM Tris–HCl, pH 8; 10 mM EDTA; 1% SDS was added to the beads for reverse crosslinking at 65C overnight with shaking. After reverse crosslinking, the beads were removed. The eluted DNA was treated with RNaseA and Proteinase K and then purified using Qiagen MinElute PCR Purification Kit, following the manufacturer’s instructions. qPCR reactions were set up using SsoAdvanced Universal SYBR Green or SsoFast™ EvaGreen Supermix (BioRad). Reactions were run on the StepOne Plus Real-Time PCR System (Applied Biosystems) or the QuantStudio 6 Flex (Thermo Fisher). The Ct values were analyzed by the relative fraction of input method.

### Next-Generation Sequencing (NGS) amplicon preparation and analysis

To confirm homozygosity/heterozygosity of the *HBD* clones, samples’ gDNA extracted with QuickExtract DNA Kit was first amplified by PCR. The primers were designed specifically for NGS, spanning a region of < 200 bp (including the primers sequences) in which the cutsite is asymmetrically placed (e.g., 30-80 bp from the forward or the reverse primer) to capture the edited region. Subsequently, two stubber sequences are added, one for the forward primer (5’−CTTTCCCTACACGACGCTCTTCCGATCT −3’) and one for the reverse (5’-GGAGTTCAGAC GTGTGCTCTTCCGATCT −3’). After running the first PCR to amplify the genetic region of interest, the overhanging stubber sequences are used to run a second PCR with indexing primers (forward and reverse primers are premixed at 5 M). Samples are pooled and purified with SPRIselect beads, 5 mL (Beckman Coulter, B23317) using a DynaMag™-2 Magnet magnetic stand (Thermo Fisher Scientific, 12321D). NGS is performed by the Genome Engineering and Measurement Lab (GEML, ETH Zürich) using a NovaSeq 6000 Sequencing System (Illumina, 20012850) or MiSeq. Sequencing mode used was either 100 or 150 PE (paired end). Editing efficiencies were determined using CRISPResso^31^.

---

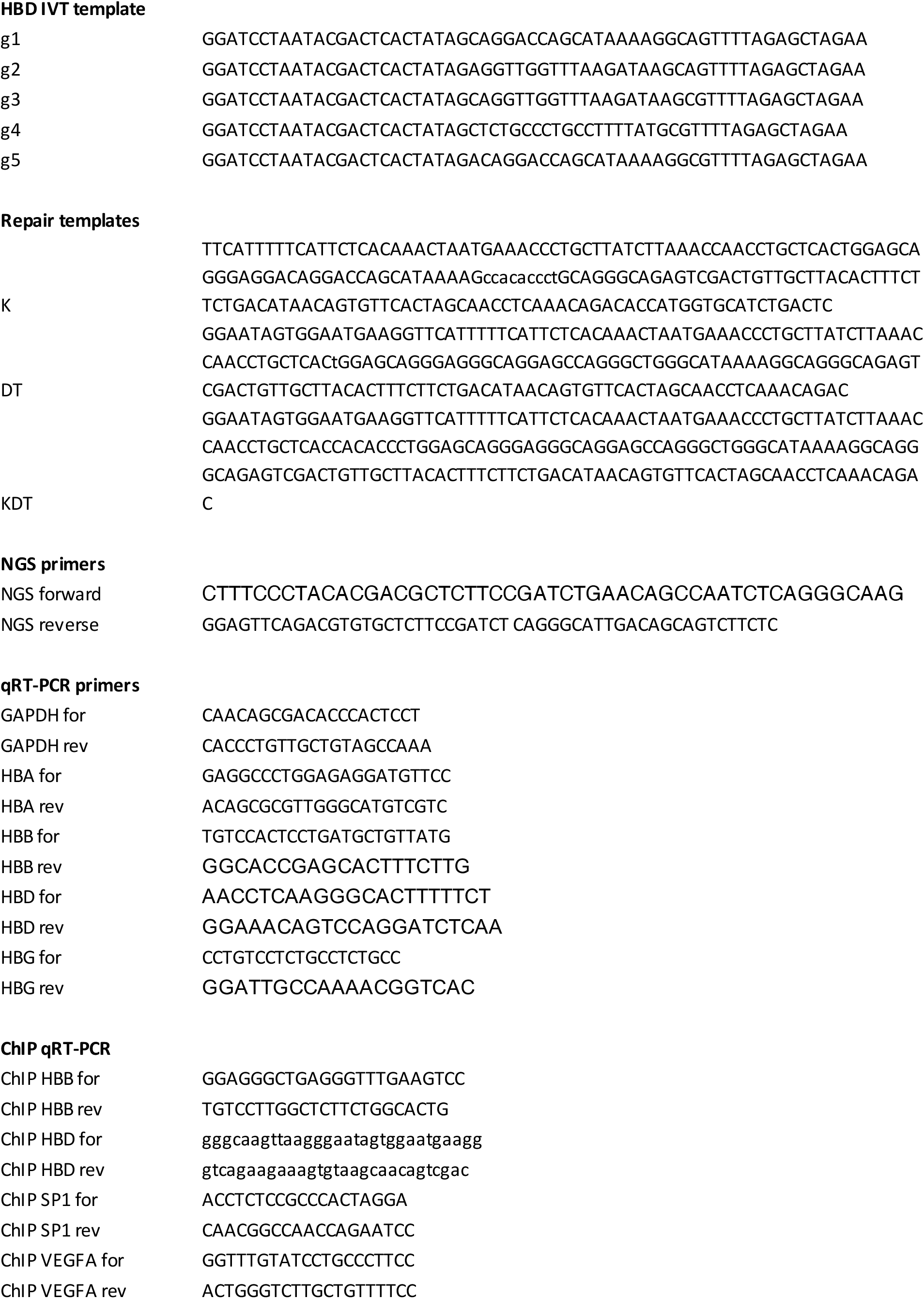

